# A pipeline approach to single-particle processing in RELION

**DOI:** 10.1101/078352

**Authors:** Rafael Fernandez-Leiro, Sjors H.W. Scheres

## Abstract

We describe the formal concept of a workflow to single-particle analysis of cryo-electron microscopy (cryo-EM) images in the RELION program. In this approach, the structure determination process is considered as a graph, where intermediate results in the form of images or metadata are the vertices, and different functionalities of the program are the edges. The new implementation automatically logs all user actions, facilitates file management and disk cleaning, and allows convenient browsing of a project’s history. Moreover, new functionality to iteratively execute consecutive jobs allows on-the-fly image processing, which will lead to more efficient data acquisition by providing faster feedback on data quality. The possibility to exchange data processing procedures among users will contribute to the development of standardised image processing procedures, and hence increase accessibility for new users in this rapidly expanding field.

## 1 Introduction

Recent advances in cryo-EM instrumentation and image processing software have substantially expanded the scope of structure determination by single-particle analysis (Fernandez-Leiro and Scheres, 2016). As a result, the field is growing fast. With many new users turning to cryo-EM structure determination, efforts in methods development are increasingly focused on improving the accessibility of the technique. Examples of this are the development of robots for sample preparation and transfer of the sample into the microscope (Cheng et al., 2007; Vos et al., 2008; Kim et al., 2010; Coudray et al., 2011), the introduction of automated data acquisition and processing software (Suloway et al., 2005; Stagg et al., 2006; Mastronarde, 2005; Li et al., 2015), the introduction of image processing software suites with convenient graphical user interfaces (GUIs) (Tang et al., 2007; Hohn et al., 2007; Baxter et al., 2007; de la Rosa-Trevin et al., 2013), and the development of integrated software environments and file formats that allow convenient interchanging between the different programs (Lander et al., 2009; de la Rosa-Trevin et al., 2016; Marabini et al., 2016). It is interesting to note that a similar development towards automation and ease-of-use happened in the related field of macromolecular structure determination by X-ray crystallography during the 1990s and 2000s (Blundell et al., 2002).

This paper describes the implementation of a pipeline approach to cryo-EM structure determination protocols in the RELION program (Scheres, 2012b). RELION is based around an empirical Bayesian approach to single-particle analysis (Scheres, 2012a), where important parameters that describe the signal-to-noise ratios of the data are inferred from the data themselves. This reduces the need for expertise to run the program, and makes it intrinsically suitable for automation. Since its introduction in 2012, a convenient GUI has further enriched the ease-of-use of RELION, but until now the concept of a workflow did not exist explicitly within the program. To facilitate the generation of standardized and (semi-)automated procedures for structure determination, we describe here the formal description of a workflow, which we call a pipeline, in the latest RELION release (version 2.0).

## 2 Implementation of the pipeline

### 2.1 Jobs and nodes

The process of cryo-EM structure determination can be described as a directed acyclic graph, consisting of vertices and edges. We will refer to the vertices as *nodes*, and a total of 14 types of different types of nodes, representing different forms of data or metadata, have been defined in RELION-2.0 (Table 6). Depending on the type, nodes are stored on the computer disk in the form of STAR (Hall, 1991), PDF, MRC (Cheng et al., 2015) or plain text files. The edges of the graph represent processes that act on these data. The edges are called *jobs*, and a total of 18 different types of job have been implemented (Table 6). With the exception of the ‘Import’ job, all job types take one or more nodes as input, and produce one or more nodes as output. Different job types take different types of input and output nodes (see Table 6). Using output nodes from one job as input for the next job builds up a graph, which we refer to as the *pipeline*.

**Table 1:**
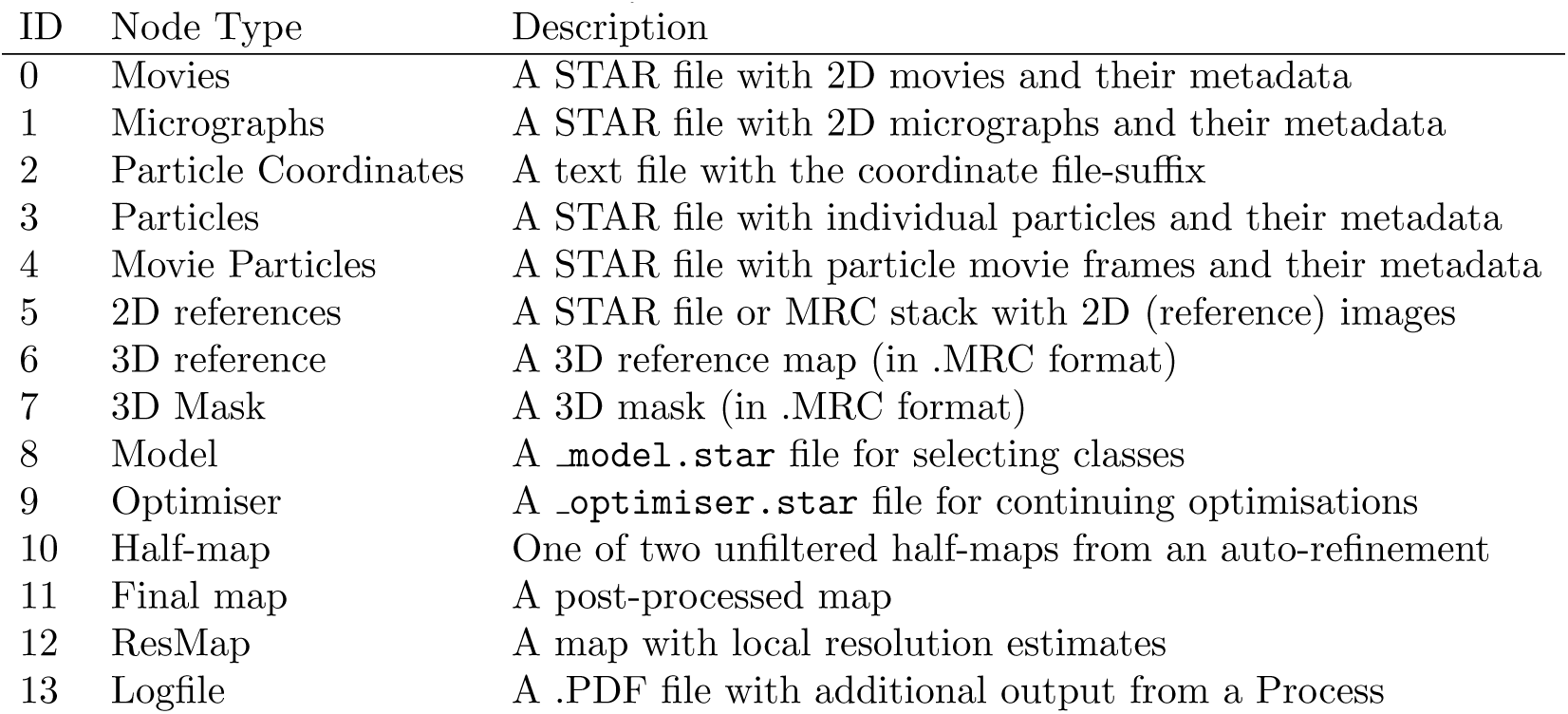
Node types in RELION-2.0

| ID | Node Type | Description |
| --- | --- | --- |
| 0 | Movies | A STAR file with 2D movies and their metadata |
| 1 | Micrographs | A STAR file with 2D micrographs and their metadata |
| 2 | Particle Coordinates | A text file with the coordinate file-suffix |
| 3 | Particles | A STAR file with individual particles and their metadata |
| 4 | Movie Particles | A STAR file with particle movie frames and their metadata |
| 5 | 2D references | A STAR file or MRC stack with 2D (reference) images |
| 6 | 3D reference | A 3D reference map (in .MRC format) |
| 7 | 3D Mask | A 3D mask (in .MRC format) |
| 8 | Model | A `_model.star` file for selecting classes |
| 9 | Optimiser | A `_optimiser.star` file for continuing optimisations |
| 10 | Half-map | One of two unfiltered half-maps from an auto-refinement |
| 11 | Final map | A post-processed map |
| 12 | ResMap | A map with local resolution estimates |
| 13 | Logfile | A .PDF file with additional output from a Process |

**Table 2:**
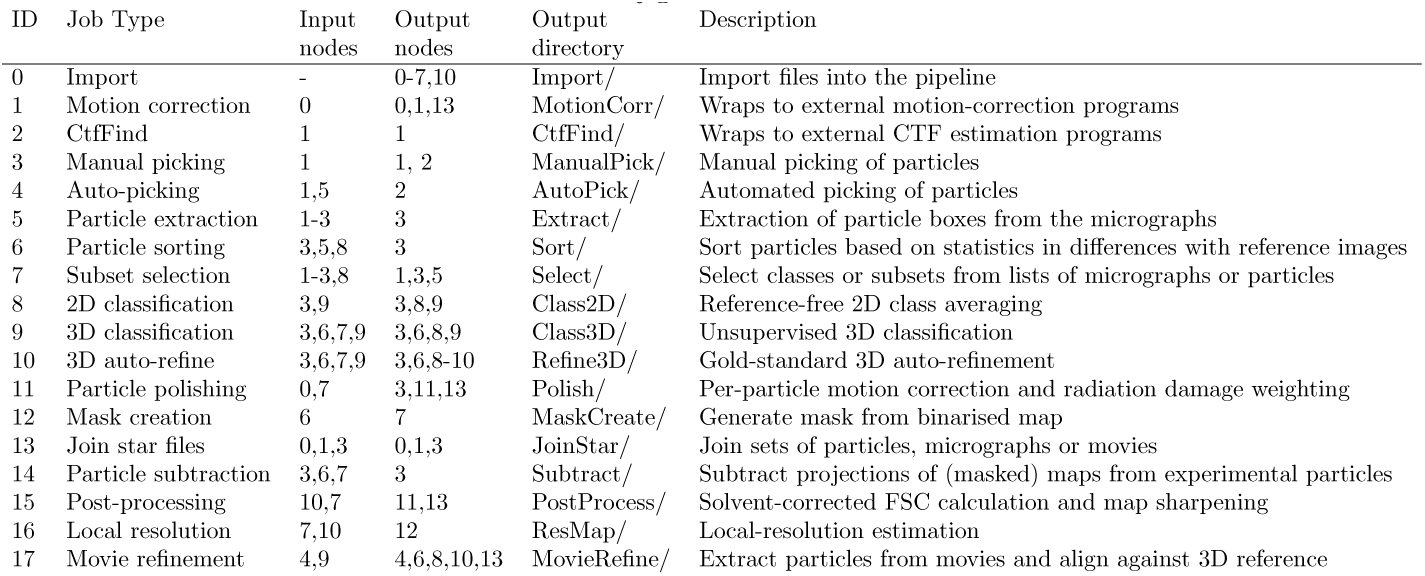
Job types in RELION-2.0.

| ID | Job Type | Input nodes | Output nodes | Output directory | Description |
| --- | --- | --- | --- | --- | --- |
| 0 | Import | - | 0-7,10 | Import/ | Import files into the pipeline |
| 1 | Motion correction | 0 | 0,1,13 | MotionCorr/ | Wraps to external motion-correction programs |
| 2 | CtfFind | 1 | 1 | CtfFind/ | Wraps to external CTF estimation programs |
| 3 | Manual picking | 1 | 1, 2 | ManualPick/ | Manual picking of particles |
| 4 | Auto-picking | 1,5 | 2 | AutoPick/ | Automated picking of particles |
| 5 | Particle extraction | 1-3 | 3 | Extract/ | Extraction of particle boxes from the micrographs |
| 6 | Particle sorting | 3,5,8 | 3 | Sort/ | Sort particles based on statistics in differences with reference images |
| 7 | Subset selection | 1-3,8 | 1,3,5 | Select/ | Select classes or subsets from lists of micrographs or particles |
| 8 | 2D classification | 3,9 | 3,8,9 | Class2D/ | Reference-free 2D class averaging |
| 9 | 3D classification | 3,6,7,9 | 3,6,8,9 | Class3D/ | Unsupervised 3D classification |
| 10 | 3D auto-refine | 3,6,7,9 | 3,6,8-10 | Refine3D/ | Gold-standard 3D auto-refinement |
| 11 | Particle polishing | 0,7 | 3,11,13 | Polish/ | Per-particle motion correction and radiation damage weighting |
| 12 | Mask creation | 6 | 7 | MaskCreate/ | Generate mask from binarised map |
| 13 | Join star files | 0,1,3 | 0,1,3 | JoinStar/ | Join sets of particles, micrographs or movies |
| 14 | Particle subtraction | 3,6,7 | 3 | Subtract/ | Subtract projections of (masked) maps from experimental particles |
| 15 | Post-processing | 10,7 | 11,13 | PostProcess/ | Solvent-corrected FSC calculation and map sharpening |
| 16 | Local resolution | 7,10 | 12 | ResMap/ | Local-resolution estimation |
| 17 | Movie refinement | 4,9 | 4,6,8,10,13 | MovieRefine/ | Extract particles from movies and align against 3D reference |

The 18 job types have all been implemented in a new GUI (Figure 1), which aims to encapsulate the entire functionality that is needed to perform cryo-EM structure determination. One can start with the import of a set of 2D micrographs or movies, and proceed from there. (Note however that *de novo* generation of a 3D reference model from the experimental images has not been implemented, and 3D references generated in other programs need to be imported into the pipeline.) New jobs can be selected from the job-type list on the top left of the GUI. Selecting a job-type from this list will load the corresponding parameter input fields on the top right part of the GUI. In order to reduce the risk of selecting incorrect input files, the fields that correspond to input nodes have ‘Browse’ buttons that will only display nodes with the expected type that already exists in the pipeline. Internally, the node-type-specific browse buttons use a hidden directory called `.Nodes/`. Sometimes, this directory gets corrupted, in which case it can be re-generated using the ‘Remake .Nodes/’ entry from the File menu on the GUI.

**Figure 1:**
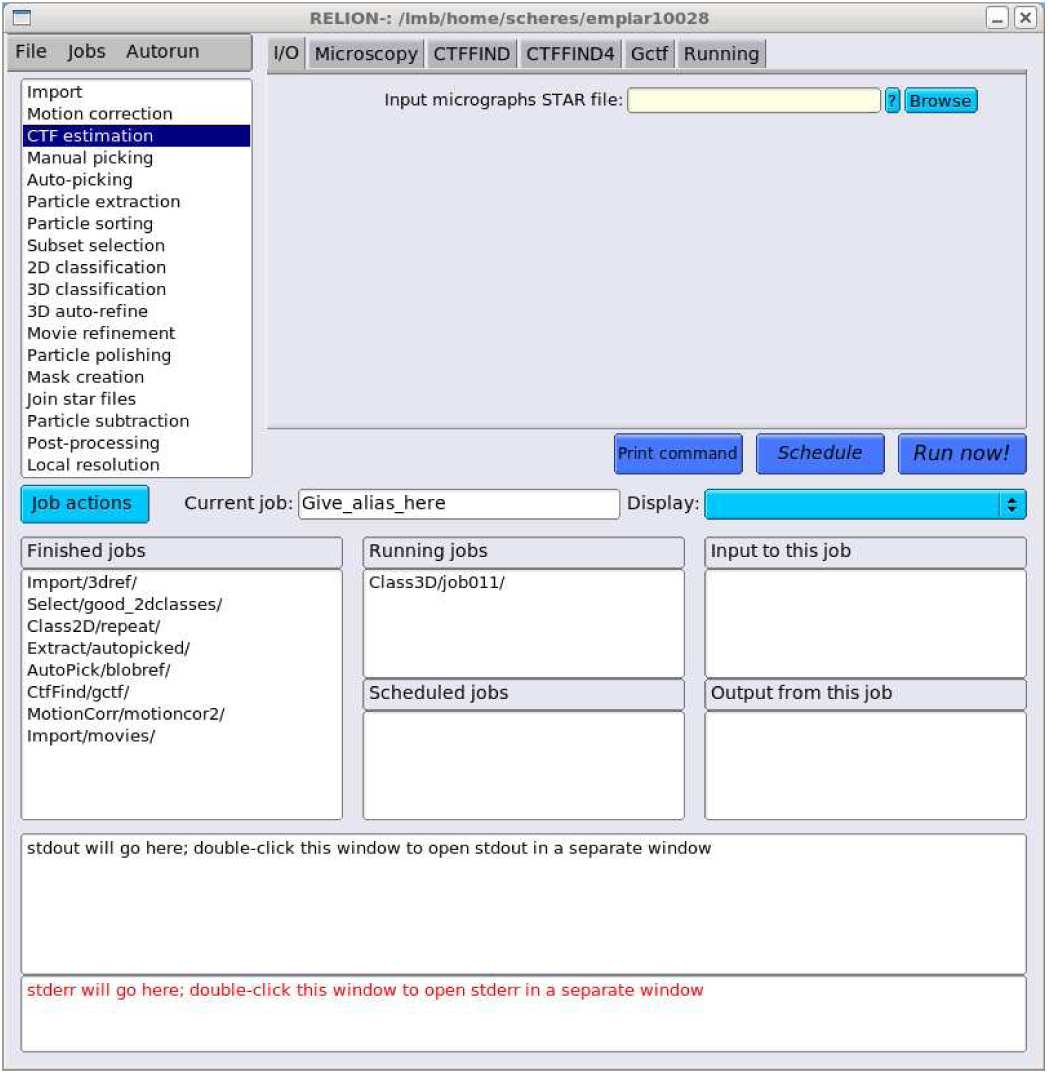
The RELION-2.0 GUI

### 2.2 Job execution

When all parameters of a job have been selected, a job can be executed using the ‘Run now!’ button on the GUI. All jobs that are executed through the GUI will be added to the pipeline, which is stored as a STAR file on the computer disk. This STAR file contains lists of all nodes and all processes in the pipeline, as well as lists of which nodes are used as input or output for which jobs. All jobs within a pipeline are numbered consecutively, and a new directory is created inside the project directory for every new job that is executed. These directories are nested inside a higher directory that reflects the job type. For example, if a 2D classification job is the tenth job to be executed in the pipeline, then its output will be stored in a directory called `Class2D/job010/`. (See Table 6 for the higher directory names for all job types). To facilitate the recognition of jobs by the user, the concept of a job alias has also been implemented. An alias is a unique name, a text string that does not contain special characters like spaces or slashes, that provides a more intuitive description for a job. The creation of an alias to a job leads to the generation of a symbolic link to that job’s output directory on the computer disk.

Upon execution of the job, its name (either its output directory or its alias) is added to a list of ‘Running jobs’ on the GUI, and it will remain there until the files of all output nodes have been detected on the computer disk. When the files of all expected output nodes are present, the status of the job will change from running to finished, and on the GUI the job will move to the list of ‘Finished jobs’. When a job is executed, the GUI also creates a text file called `note.txt` inside the job’s output directory, which will contain a time stamp when the job was executed and the exact command line arguments of the program used. The GUI will also open a text editor window that allows the user to further comment on the intentions or characteristics of the job. The latter could serve as an electronic logbook for the user.

Some jobs are stopped before they reach their intended result. In order to finish such jobs, the ‘Run now!’ button on the GUI changes to a ‘Continue now’ button when a job is selected from any of the lists in the lower part of the GUI. Continuing a job works in different manners for different types of jobs. Pre-processing jobs like ‘Motion correction’, ‘CTF estimation’, ‘Manual picking’, ‘Auto-picking’ and ‘Particle extraction’, as well as ‘Movie refinement’ will skip any micrographs for which the expected output files are already on the computer disk. The likelihood optimisation jobs ‘2D classifcation’, ‘3D classification’ and ‘3D auto-refine’ can be continued from their last finished iteration by providing the corresponding `_optimiser.star` file. Sometimes, it is not necessary to continue a job that was stopped prematurely. In that case, the user can also use the ‘Mark as finished’ option under the ‘Job action’ button, which will move the job from the running to the finished job lists. In case of a likelihood optimisation job, the expected output nodes in the pipeline will then be replaced for the output files of the last finished iteration.

It is important to note that RELION does not retain control over the running job: it will merely wait for the expected output files to appear on the computer disk. This means that there is no functionality for the user to kill a submitted job through the GUI, which should instead be done through the operating system or the job queue.

### 2.3 Browsing the project history

The user can conveniently browse through all jobs in the pipeline by clicking jobs in the lists of ‘Finished jobs’, ‘Running jobs’ or ‘Scheduled jobs’. When the users clicks on a job in these lists, the parameters of that job get read from a file called run.job inside that job’s output directory, and these get loaded into the parameter input fields on the top part of the GUI. Moreover, any (upstream) jobs that generated the input nodes for the selected job are shown on the GUI in the list called ‘Input to this job’, and any (downstream) jobs that take the output nodes of this job as input are shown in the list called ‘Output from this job’. Clicking on jobs in the latter two lists allows convenient browsing back and forth through the history of the project. In addition, the user can explore how jobs are connected to each other by generating vector-based, and thereby conveniently editable, flowcharts in PDF format.

### 2.4 Analysing results

For each job in the pipeline, the ‘Display’ button on the GUI allows visualisation of its own input and output nodes. This guides the user by presenting only a few options of which files to display for each job. For several job types, it is often worthwhile to also inspect part of the intermediate results. To this purpose, some job types will output a file called `logfile.pdf`, which presents the most relevant necessary intermediate results in a convenient, graphical manner. Examples include the movement tracks of individual movies from ‘Motion correction’ jobs; the B-factor plots and per-particle movement tracks from ‘Particle polishing’; and FSC as well as Guinier plots from ‘Post-processing’ jobs.

A more general display functionality that allows visualisation of any image, reminiscent of the ‘Display’ button in previous versions of RELION, is still available through the ‘File’ menu.

### 2.5 Disk management

In a typical structure determination project, pipelines can quickly become complicated, as many different ways to process their data are explored. This may generate large amounts of intermediate results that occupy substantial amounts of space on the computer disk. The strict organisation of output files in separate directories for each job facilitates managing these files to limit disk space. The GUI implements functionality to delete jobs that are deemed to be no longer necessary by removing their output directories from the project directory. In addition, the user can also choose to clean output directories of jobs, in which case only intermediate files are removed, but files necessary for the pipeline are retained. Both functionalities are accessible through the ‘Job actions’ button. To protect the user from unwanted loss of data, upon deletion or cleaning of a job, all output files are initially moved to a `Trash/` folder inside the project directory. Entire jobs can be ‘undeleted’ from the ‘Jobs’ menu, while specific files can also be recovered manually using the linux command line. To free disk space, files can be removed permanently through an ‘Empty Trash’ option on the ‘File’ menu.

## 3 On-the-fly processing and exchanging procedures

Instead of immediately executing a job once all its parameters have been selected, the user can also use the ‘Schedule’ button to schedule a job for execution at a later stage. Even though the files for the output nodes of scheduled jobs do not exist yet, they will still be added to the `.Nodes/` directory that is used by the ‘Browse’ buttons on the parameter input fields. Thereby, once a job is scheduled, one can select its expected output nodes as input for another job, which can then also be scheduled. In this manner, one can schedule multiple, consecutive jobs for future execution.

Execution of scheduled jobs can be performed through the ‘Autorun’ menu on the GUI. This will launch a separate window, in which the user can select which of the scheduled jobs to execute. In addition, the user can opt to cycle through the selected scheduled jobs multiple times, and specify a minimum time between subsequent iterations. Combined with the functionality to continue jobs explained above, this provides a simple mechanism for on-the-fly processing of data during acquisition. For example, one could schedule an ‘Import’ job that imports all movie files in a given directory, followed by ‘Motion correction’, ‘CTF estimation’, ‘Auto-picking’, and ‘Particle extraction’ jobs. This cycle of consecutive jobs could be repeated many times while the data are being acquired. In each iteration, the jobs will only act on those movies that had not been processed before.

Once enough particles have been extracted, the output from the ‘Particle extraction’ job above may also be used as input for a ‘2D classification’ job. This will lead to the calculation of reference-free 2D class averages, which typically provide useful insights into the quality of the data. Probably one would not want to perform the (computationally more expensive) 2D classification job as often as one would want to pre-process the incoming movies. Therefore, multiple executions of scheduled jobs can be run independently. For example, one could pre-process new micrographs from ‘Import’ to ‘Particle extraction’ every 5 minutes, but only execute ‘2D classification’ with the extracted particles every hour. Provided sufficient computer resources are at hand to process the incoming data, this procedure allows the user to assess data quality from the inspection of 2D class averages during data acquisition. Thereby, on-the-fly data processing will allow the user to change his data acquisition scheme in case the data is unsatisfactory. Even ‘3D classification’ or ‘3D auto-refine’ jobs may be included in on-the-fly processing. This may be particularly useful to assess whether co-factors are bound to complexes, or whether the acquired data is capable of reaching high-resolution.

The capability of executing multiple, previously scheduled jobs is also relevant for developing standardised procedures for image processing. To facilitate this, scheduled jobs can be exported from the pipeline using the corresponding option from the ‘Jobs’ menu. Exporting scheduled jobs will change the directory structure with the numbered jobs from the current pipeline to a more generic directory structure. This generic structure can then be copied into the directory of a different project, where it can be imported into the pipeline by again using the corresponding option from the ‘Jobs’ menu. This will allow different users to share their image processing procedures, which will be of help for inexperienced users and may further facilitate the development of automated and standardised procedures.

## 4 Results

### 4.1 Test case description

To demonstrate the new pipeline, we reprocessed a data set from the EMPIAR data base (Iudin et al., 2016): entry 10028 (Wong et al., 2014). This cryo-EM data set comprises 1,081 16-frame movies of 4,096 × 4,096 pixels that were collected using a Falcon-II direct-electron detector on an FEI Polara microscope. The sample contained a mixture of (cytoplasmic) 80S ribosomes (at 160 nM) that were purified from *Plasmodium falciparum* parasites, and a 1 mM solution of the anti-protozoan drug emetine. In the original study, the structure was solved to an overall resolution of 3.2 *Å* using a beta-version of RELION-1.3.

All experiments described below were performed on a single desktop machine, equipped with a Intel i7-6800K 3.4 GHz 6-core CPU, 64 Gb or RAM, and two Titan-X (Pascal) GPUs, which was recently acquired for less than £3,000. GPU-acceleration was used for motion-correction in MotionCor2 (Zheng et al., 2016), CTF parameter estimation in Gctf (Zhang, 2016), as well as for auto-picking, classification and refinement in RELION-2.0 (Kimanius et al., 2016).

### 4.2 Simulated on-the-fly-processing

In an attempt to simulate the data acquisition process, we copied a single movie per minute into a micrographs directory. The copying process was started at 14:00 in the afternoon and continued throughout the night until 08:30 the morning after. Although this is admittedly an over-simplification of the data acquisition process (for example the data set only contains high-quality images, did not stall, *etc*), our simulation still illustrates the functionality of on-the-fly data processing, which is already in active use at our cryo-EM facility.

To perform on-the-fly data processing, we scheduled consecutive jobs to *(i)* import all movies in the micrographs directory; *(ii)* run MotionCor2; *(iii)* run Gctf; *(iv)* run auto-picking; and *(v)* extract the particles. The scheduled jobs were then executed in a continuous loop, with a minimum of one minute between subsequent iterations, during the entire copying process. Motion correction was done in five patches and included dose-weighting. Gctf estimation used equiphase averaging to increase the signal-to-noise ratios in the Thon-ring images. Auto-picking (Scheres, 2015) was performed with a single Gaussian blob as template. The latter is a new option in RELION-2.0 and involves the choice of two parameters: the width and the height of the Gaussian. The width of the Gaussian was chosen to be similar to the expected diameter of the particles (270 Å). The height of the Gaussian is affected by the signal-to-noise ratio in the images, and a suitable value (0.3) was determined by a test run on the first few micrographs. Three-fold downscaled particles were initially extracted in boxes of 120 × 120 pixels. The two GPUs on our machine were used to process two movies in parallel, if needed. The pre-processing procedure took approximately one minutes per movie, half of which was taken by MotionCor2. Consequently, data processing could be done as fast as the data were coming in, and the entire pre-processing finished only four minutes after the last movie was copied.

After 111 movies had been copied, a second independent loop of ‘2D classification’ jobs was also executed, with a minimum of one hour between subsequent iterations. These jobs generated reference-free 2D class averages for all of the extracted particles that had been acquired thus far. These jobs competed for the same two GPUs that were also used by the pre-processing jobs. The execution times for the 2D classification jobs gradually increased as more particles were extracted from the available minutes. The first job took only 50 minutes, whereas the last job took 3.5 hours and finished at 12:30 on the next day, i.e. four hours after the data copying ended. Figure 2A shows how the class averages developed throughout the copying process. Excellent 2D class averages, showing different projections of 80S ribosomes with many protein and RNA-like features, confirmed the high quality of the data set already during the early stages of copying.

**Figure 2:**
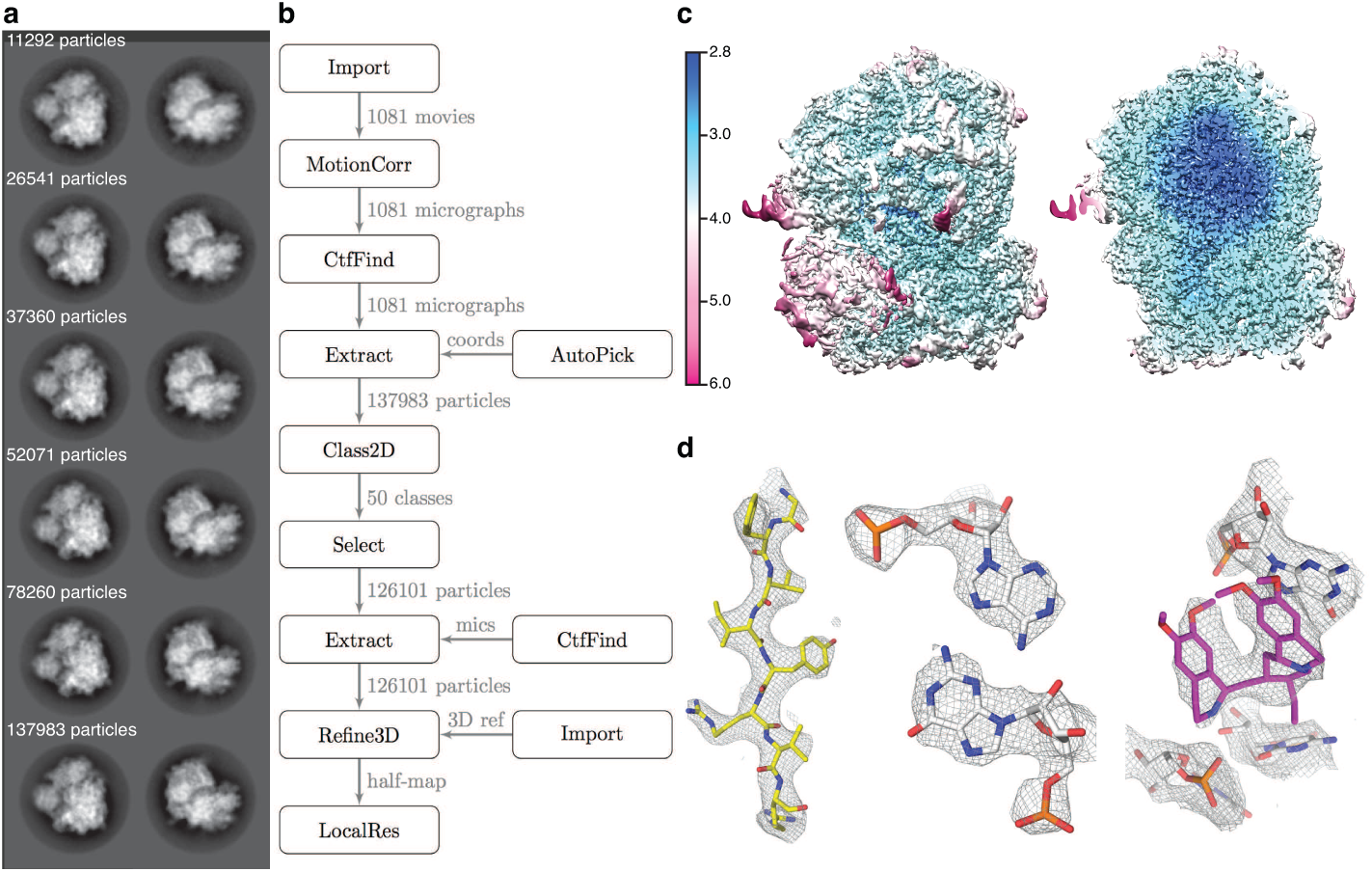
Results for the EMPIAR-10028 data set. A. Two representative reference-free 2D class averages at different times during the data copying process. B. Automatically generated flowchart of the data processing process. C. Front view and slice through a map that is filtered according to the local resolution as estimated by RELION. D. Close-up views of details in the map (a beta-strand, a RNA base pair, and the emetine compound).

Subsequently, by manually inspecting the final 2D class averages, we selected 126,101 particles, re-extracted these in boxes of 360 × 360 pixels (without down-scaling), and used these as input for a ‘3D auto-refine’ job. We used the same initial 3D reference (EMDB-2275 (Bai et al., 2013)) as was used in the original study (Wong et al., 2014). The refinement was followed by standard ‘Mask creation’ and ‘Post-processing’ jobs. Finally, to estimate the variations in local resolution throughout the reconstruction, we used a new feature in the ‘Local resolution’ job of RELION-2.0. In this feature, we use phase-randomisation (Chen et al., 2013) to correct for the solvent-effects of a small, spherical mask, which is slided over the entire map. In total, 3D auto-refine, post-processing and local resolution estimation took 16 hours on our desktop machine. Figure 2B shows the automatically generated flowchart of the data processing pipeline; figure 2C shows the local resolution estimates; and figure 2D shows details of the reconstructed density on top of the original atomic model (Wong et al., 2014).

### 4.3 Standardisation and procedure exchange

To illustrate the functionality to exchange data processing procedures among users, we exported the entire pipeline for the re-processing of the EMPIAR10028 data set, as described above, and submitted a compressed TAR archive as supplementary information to this paper. Users interested in adopting or adapting this procedure can uncompress this archive in their project directory and import the file called `./export1/exported.star` through the corresponding option on the ‘Jobs’ menu.

## 5 Conclusion

We describe how a pipelined approach to cryo-EM structure determination formalises the concept of a directed acyclic graph in RELION-2.0. The new approach improves the user experience by reducing the scope for errors, automating bookkeeping, standardising the organisation of output files, allowing straightforward browsing through the project’s history, and convenient disk cleaning functionalities. The functionality to schedule multiple, consecutive jobs for future execution, and the option to execute these jobs in iterative loops allows on-the-fly processing of the data while it is being acquired. This will lead to more efficient use of microscopy time, as more informed assessments of data quality can be made while that data is still being acquired. Finally, the possibility to export and import (parts of) pipelines will allow the exchange of data processing strategies between users, which will further improve the accessibility of the technique to inexperienced users, and facilitate the development of standardised data processing procedures. RELION-2.0 is open-source software. For download instructions and documentation, please refer to the RELION Wiki pages at www2.mrc-lmb.cam.ac.uk/relion. For user questions and feedback, please use the CCPEM mailing list at www.jiscmail.ac.uk/CCPEM.

## Acknowledgements

This work was funded by the UK Medical Research Council (MC UP A025 1013 to SHWS).

## References

Bai, X.-C., Fernandez, I. S., McMullan, G., Scheres, S. H., 2013. Ribosome structures to near-atomic resolution from thirty thousand cryo-EM particles. eLife 2, e00461.

Baxter, W. T., Leith, A., Frank, J., Jan. 2007. SPIRE: the SPIDER reconstruction engine. Journal of Structural Biology 157 (1), 56–63.

Blundell, T. L., Jhoti, H., Abell, C., Jan. 2002. High-throughput crystallography for lead discovery in drug design. Nature Reviews. Drug Discovery 1 (1), 45–54.

Chen, S., McMullan, G., Faruqi, A. R., Murshudov, G. N., Short, J. M., Scheres, S. H. W., Henderson, R., Dec. 2013. High-resolution noise substitution to measure overfitting and validate resolution in 3d structure determination by single particle electron cryomicroscopy. Ultramicroscopy 135, 24–35.

Cheng, A., Henderson, R., Mastronarde, D., Ludtke, S. J., Schoenmakers, R. H. M., Short, J., Marabini, R., Dallakyan, S., Agard, D., Winn, M., Nov. 2015. MRC2014: Extensions to the MRC format header for electron cryo-microscopy and tomography. Journal of Structural Biology 192 (2), 146–150.

Cheng, A., Leung, A., Fellmann, D., Quispe, J., Suloway, C., Pulokas, J., Abeyrathne, P. D., Lam, J. S., Carragher, B., Potter, C. S., Dec. 2007. Towards automated screening of two-dimensional crystals. Journal of Structural Biology 160 (3), 324–331.

Coudray, N., Hermann, G., Caujolle-Bert, D., Karathanou, A., Erne-Brand, F., Buessler, J.-L., Daum, P., Plitzko, J. M., Chami, M., Mueller, U., Kihl, H., Urban, J.-P., Engel, A., Rmigy, H.-W., Feb. 2011. Automated screening of 2d crystallization trials using transmission electron microscopy: a high-throughput tool-chain for sample preparation and microscopic analysis. Journal of Structural Biology 173 (2), 365–374.

de la Rosa-Trevin, J. M., Otn, J., Marabini, R., Zaldvar, A., Vargas, J., Carazo, J. M., Sorzano, C. O. S., Nov. 2013. Xmipp 3.0: an improved software suite for image processing in electron microscopy. Journal of Structural Biology 184 (2), 321–328.

de la Rosa-Trevin, J. M., Quintana, A., Del Cano, L., Zaldvar, A., Foche, I., Gutirrez, J., Gmez-Blanco, J., Burguet-Castell, J., Cuenca-Alba, J., Abrishami, V., Vargas, J., Otn, J., Sharov, G., Vilas, J. L., Navas, J., Conesa, P., Kazemi, M., Marabini, R., Sorzano, C. O. S., Carazo, J. M., Jul. 2016. Scipion: A software framework toward integration, reproducibility and validation in 3d electron microscopy. Journal of Structural Biology 195 (1), 93–99.

Fernandez-Leiro, R., Scheres, S. H. W., 2016. Unravelling biological macro-molecules with cryo-electron microscopy. Nature 537 (7620), 339–346.

Hall, S. R., 1991. The STAR file: a new format for electronic data transfer and archiving. J. Chem. Inf. Comput. Sci. 31 (2), 326–333.

Hohn, M., Tang, G., Goodyear, G., Baldwin, P. R., Huang, Z., Penczek, P. A., Yang, C., Glaeser, R. M., Adams, P. D., Ludtke, S. J., Jan. 2007. SPARX, a new environment for Cryo-EM image processing. Journal of Structural Biology 157 (1), 47–55.

Iudin, A., Korir, P. K., Salavert-Torres, J., Kleywegt, G. J., Patwardhan, A., May 2016. EMPIAR: a public archive for raw electron microscopy image data. Nature Methods 13 (5), 387–388.

Kim, C., Vink, M., Hu, M., Love, J., Stokes, D. L., Ubarretxena-Belandia, I., Jun. 2010. An automated pipeline to screen membrane protein 2d crystallization. Journal of Structural and Functional Genomics 11 (2), 155–166.

Kimanius, D., Forsberg, B. O., Scheres, S., Lindahl, E., Jun. 2016. Accelerated cryo-EM structure determination with parallelisation using GPUs in RELION-2. bioRxiv, 059717.

Lander, G. C., Stagg, S. M., Voss, N. R., Cheng, A., Fellmann, D., Pulokas, J., Yoshioka, C., Irving, C., Mulder, A., Lau, P.-W., Lyumkis, D., Potter, C. S., Carragher, B., Apr. 2009. Appion: an integrated, database-driven pipeline to facilitate EM image processing. Journal of Structural Biology 166 (1), 95–102.

Li, X., Zheng, S., Agard, D. A., Cheng, Y., Nov. 2015. Asynchronous data acquisition and on-the-fly analysis of dose fractionated cryoEM images by UCSFImage. Journal of Structural Biology 192 (2), 174–178.

Marabini, R., Ludtke, S. J., Murray, S. C., Chiu, W., de la Rosa-Trevin, J. M., Patwardhan, A., Heymann, J. B., Carazo, J. M., May 2016. The Electron Microscopy eXchange (EMX) initiative. Journal of Structural Biology 194 (2), 156–163.

Mastronarde, D. N., Oct. 2005. Automated electron microscope tomography using robust prediction of specimen movements. Journal of Structural Biology 152 (1), 36–51.

Scheres, S. H. W., Jan. 2012a. A Bayesian view on cryo-EM structure determination. Journal of Molecular Biology 415 (2), 406–418.

Scheres, S. H. W., Dec. 2012b. RELION: Implementation of a Bayesian approach to cryo-EM structure determination. Journal of Structural Biology 180 (3), 519–530.

Scheres, S. H. W., Feb. 2015. Semi-automated selection of cryo-EM particles in RELION-1.3. Journal of Structural Biology 189 (2), 114–122.

Stagg, S. M., Lander, G. C., Pulokas, J., Fellmann, D., Cheng, A., Quispe, J. D., Mallick, S. P., Avila, R. M., Carragher, B., Potter, C. S., Sep. 2006. Automated cryoEM data acquisition and analysis of 284742 particles of GroEL. Journal of Structural Biology 155 (3), 470–481.

Suloway, C., Pulokas, J., Fellmann, D., Cheng, A., Guerra, F., Quispe, J., Stagg, S., Potter, C. S., Carragher, B., Jul. 2005. Automated molecular microscopy: the new Leginon system. Journal of Structural Biology 151 (1), 41–60.

Tang, G., Peng, L., Baldwin, P. R., Mann, D. S., Jiang, W., Rees, I., Ludtke, S. J., Jan. 2007. EMAN2: an extensible image processing suite for electron microscopy. Journal of Structural Biology 157 (1), 38–46.

Vos, M. R., Bomans, P. H. H., Frederik, P. M., Sommerdijk, N. A. J. M., Oct. 2008. The development of a glove-box/Vitrobot combination: Airwater interface events visualized by cryo-TEM. Ultramicroscopy 108 (11), 1478–1483.

Wong, W., Bai, X.-c., Brown, A., Fernandez, I. S., Hanssen, E., Condron, M., Tan, Y. H., Baum, J., Scheres, S. H. W., 2014. Cryo-EM structure of the Plasmodium falciparum 80s ribosome bound to the anti-protozoan drug emetine. eLife 3.

Zhang, K., Jan. 2016. Gctf: Real-time CTF determination and correction. Journal of Structural Biology 193 (1), 1–12.

Zheng, S., Palovcak, E., Armache, J.-P., Cheng, Y., Agard, D., Jul. 2016. Anisotropic Correction of Beam-induced Motion for Improved Single-particle Electron Cryo-microscopy. bioRxiv, 061960.

